# Clustering of desmosomal cadherins by desmoplakin is essential for cell-cell adhesion

**DOI:** 10.1101/2020.04.06.027185

**Authors:** Marie-Therès Wanuske, Dominique Brantschen, Camilla Schinner, Chiara Stüdle, Elias Walter, Matthias Hiermaier, Franziska Vielmuth, Jens Waschke, Volker Spindler

**Affiliations:** Department of Biomedicine, University of Basel, Basel, Switzerland; Faculty of Medicine, Ludwig-Maximilians-Universität Munich, Munich, Germany

## Abstract

Desmoplakin (Dp) localizes to desmosomes, linking clusters of desmosomal adhesion molecules to the intermediate filament cytoskeleton. Here, we generated Dp knockout (ko) cell lines of human keratinocytes to study the impact on desmosomal adhesion molecules and desmosome turnover using atomic force microscopy and superresolution imaging. In comparison to ko of another desmosomal component, plakoglobin (Pg), loss of Dp resulted in absence of desmosomes and drastically impaired cell cohesion. In Dp ko, the desmosomal adhesion molecules desmoglein 2 (Dsg2) and desmocollin 3 (Dsc3) were redistributed into small clusters in the cell membrane with no further increase in loss of intercellular adhesion by silencing of Dsg2. This suggests that extradesmosomal cadherins do not significantly contribute to cell cohesion but rather localization within desmosomes is required. Our data outline a crucial role of Dp for both desmosomal molecule clustering and mature desmosome formation and provide novel insights into the regulation of intercellular adhesion.

## INTRODUCTION

The human epidermis functions as a protective barrier against external irritants and mechanical stress. Desmosomes, highly organized supramolecular structures, are essential in maintaining tissue integrity by providing strong intercellular adhesion. They connect adjacent cells via the desmosomal cadherins desmogleins (Dsg) 1-4 and desmocollins (Dsc) 1-3 which trans-interact in a Ca^2+^-dependent, hetero- and homophilic manner in the extracellular space (Chitaev and Troyanovsky, 1997, Delva et al., 2009, Nie et al., 2011). The armadillo protein family members plakophilin (Pkp) 1-3 and plakoglobin (Pg) link the cadherins to the most abundant desmosomal component, desmoplakin (Dp) (Desai et al., 2009, Mueller and Franke, 1982). This plaque protein exists in two major splice variants (I and II) (Green et al., 1987) which are both expressed in the epidermis (Angst et al., 1990). Serving as a key linker, Dp tethers the adhesion complex to the intermediate filament cytoskeleton via its carboxy-terminal domain (Bornslaeger et al., 1996, Kouklis et al., 1994), while the amino-terminal domain was demonstrated to cluster Pg and desmosomal cadherins (Kowalczyk et al., 1997). In addition to these desmosome-bound clusters, desmosomal cadherins are proposed to occur extradesmosomal in the membrane. As desmosomes undergo constant reorganization, this dynamic pool may be recruited to the desmosome to rapidly adapt to external influences (Nekrasova and Green, 2013, Spindler and Waschke, 2014, Windoffer et al., 2002). However, the difference between extradesmosomal and plaque-bound cadherins remains largely unclear.

Besides *in vitro* studies, in which reduced intercellular cohesion was observed by downregulation of Dp (Cabral et al., 2012, Spindler et al., 2014), various mutations in the Dp gene have been identified to cause diverse diseases involving both the heart (Yang et al., 2006) and the skin, such as arrhythmogenic cardiomyopathy and carvajal syndrome (Corrado et al., 2017), lethal acantholytic epidermolysis (Jonkman et al., 2005) and skin fragility-woolly hair syndrome (Whittock et al., 2002). Moreover, Dp null-mutant mice do not survive beyond embryonic day 6.5 and show desmosomes lacking attachment to cytokeratin filaments (Gallicano et al., 1998), underlining the importance of Dp in maintaining physiological development and desmosome stability. Available data suggest that binding properties of desmosomal cadherins are dependent on desmosomal plaque proteins. For instance, loss of keratins weakened the binding forces of single molecule Dsg3 interactions in mouse keratinocytes (Vielmuth et al., 2018). Furthermore, modulation of the desmosomal plaque through ko of Pkp1 and Pkp3 reduced the frequency of Dsg3-mediated interactions (Fuchs et al., 2019). These results suggest that connection to the cytoskeleton or the clustering within the desmosome regulate the binding properties of desmosomal cadherins. To test this concept further, we here established Dp- and Pg-deficient human keratinocytes to investigate how these two molecules regulate the function of desmosomal cadherins and, more broadly, intercellular adhesion.

## RESULTS

### Loss of Dp but not Pg leads to lack of desmosomes and reduced intercellular adhesion

To elucidate the roles of Dp and Pg in intercellular adhesion, both plaque proteins were individually knocked out in human HaCaT keratinocytes. CRISPR/Cas9 was utilized to generate monoclonal ko and respective control (ctrl) cell lines for each protein. Effective gene targeting of all clones was validated by sequencing and effects on protein levels were confirmed by immunostaining and Western blot (Supp. Fig. 1). Initially, we evaluated the effects of Dp or Pg deficiency on the expression levels of other desmosomal proteins in three ctrl and ko cell lines each. In both models, Pkp1 was significantly reduced throughout all three ko cell lines in comparison to their corresponding ctrl pool. Whereas all Dp-deficient clones showed reduced Dsc3 levels, Dp and Dsg3 were decreased in Pg ko cells (Fig. 1a). Expression of the adherens junction proteins E-cadherin (E-Cad) and β-catenin (β-Cat) remained unaltered in Dp ko. β-Cat, which is a close homolog of Pg (Ben-Ze’ev and Geiger, 1998), was slightly upregulated under loss of Pg. E-Cad levels tended to be decreased in all Pg-deficient cell lines (Supp. Fig. 1). Next, intercellular adhesion was investigated in dissociation assays by application of mechanical stress to a detached cell monolayer. Absence of Dp led to massive increase of fragment numbers, which was less pronounced in cells lacking Pg (Fig. 1b). This suggests that the two plaque proteins contribute to global cell cohesion in variable degree. To correlate these findings with morphological appearance, transmission electron microscopy was carried out (Fig. 1c). While all ctrl cell lines maintained a normal desmosomal ultrastructure, desmosomes in Pg ko cells were morphologically compromised and anchorage of intermediate filaments appeared disrupted. In contrast, desmosomes were completely absent in cells deficient for Dp. Thus, the correlation of cell-cell adhesion with the presence of desmosomes indicates the requirement of regular desmosome formation for normal keratinocyte cohesion.

**Fig. 1.**
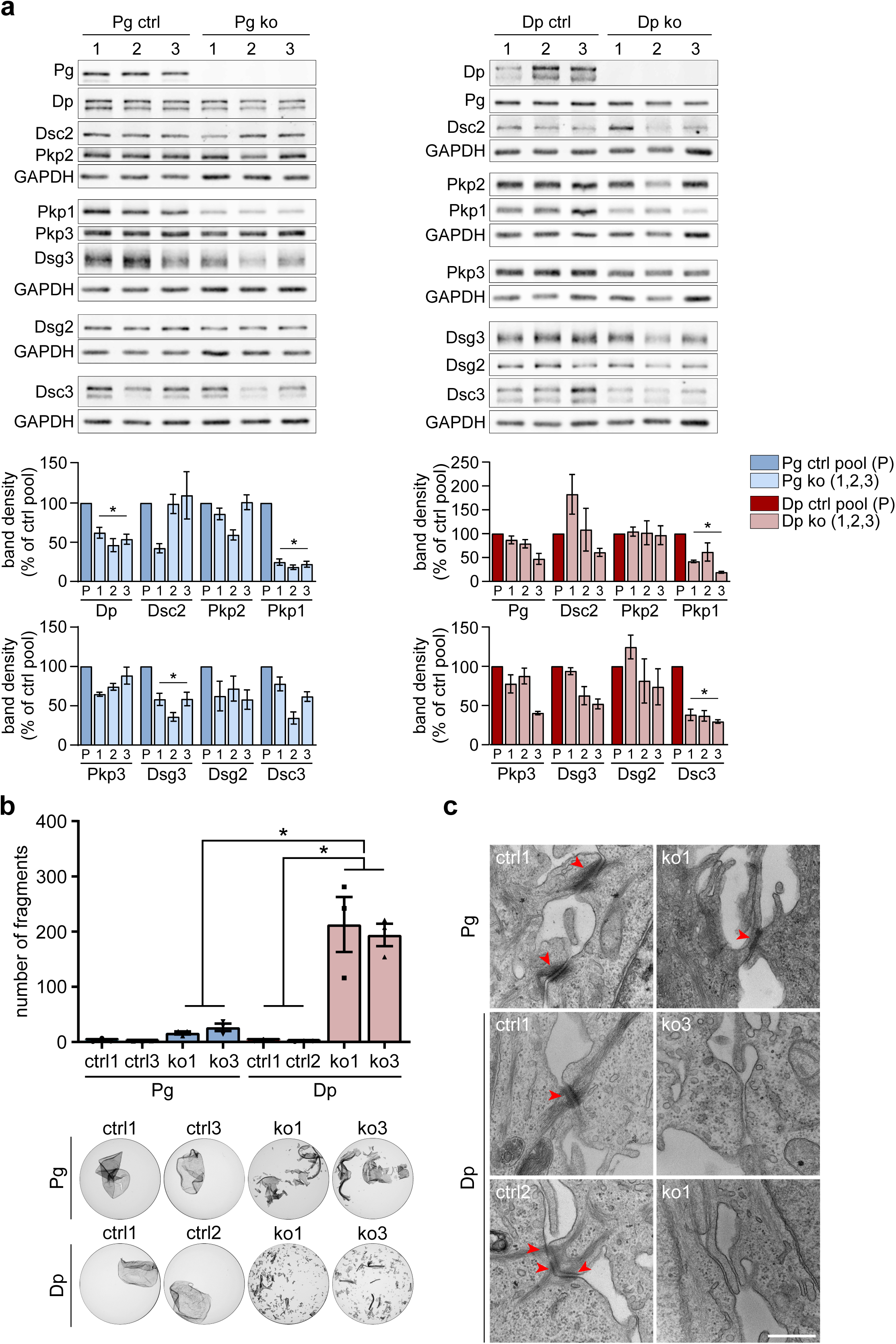
Loss of Dp but not Pg leads to lack of desmosomes and strongly impaired intercellular adhesion. (**a**) Western blot analysis of desmosomal proteins in HaCaT keratinocytes deficient for Dp or Pg and respective ctrl cell lines. GAPDH served as a loading control. Ctrl pool is the mean value of the three ctrl cell lines shown in upper panel. Each ko clone was normalized to the respective ctrl pool, n≥3, \**p*<0.05 vs. ctrl pool, error bars represent mean ± SEM. (**b**) Representative images and quantification of HaCaT monolayers after dissociation assay. Each data point represents the mean value of ≥2 replicates of one independent experiment, n=3, \**p*<0.05, error bars represent mean ± SEM. (**c**) Representative electron micrographs showing Dp and Pg ko and ctrl cell lines. Scale bar: 500 nm.

### Binding interactions of desmosomal cadherins remain unaltered under loss of Dp and Pg

Previously, we reported that binding properties of desmosomal cadherins were altered in keratin-deficient cells (Vielmuth et al., 2018), indicating that cytoskeletal anchorage affects the function of individual desmosomal cadherins. To assess how loss of Dp, the molecule mediating the anchorage to keratin filaments, influences the function of desmosomal cadherins, AFM measurements were used to examine single molecule binding properties of Dsg3 and Dsc3 as previously described (Vielmuth et al., 2015). For this purpose, AFM tips were functionalized with recombinant extracellular domains (ECD) of the respective protein. Topography and adhesion maps were generated in parallel by approaching and retracting the cantilever in a pixel-wise fashion in regions of interests both on the cell surface and at cell border areas of living keratinocyte monolayers (Fig. 2a). We confirmed specificity of Dsg3 or Dsc3 interactions in previous studies in cell free experiments (Spindler et al., 2009, Vielmuth et al., 2018).

**Fig. 2.**
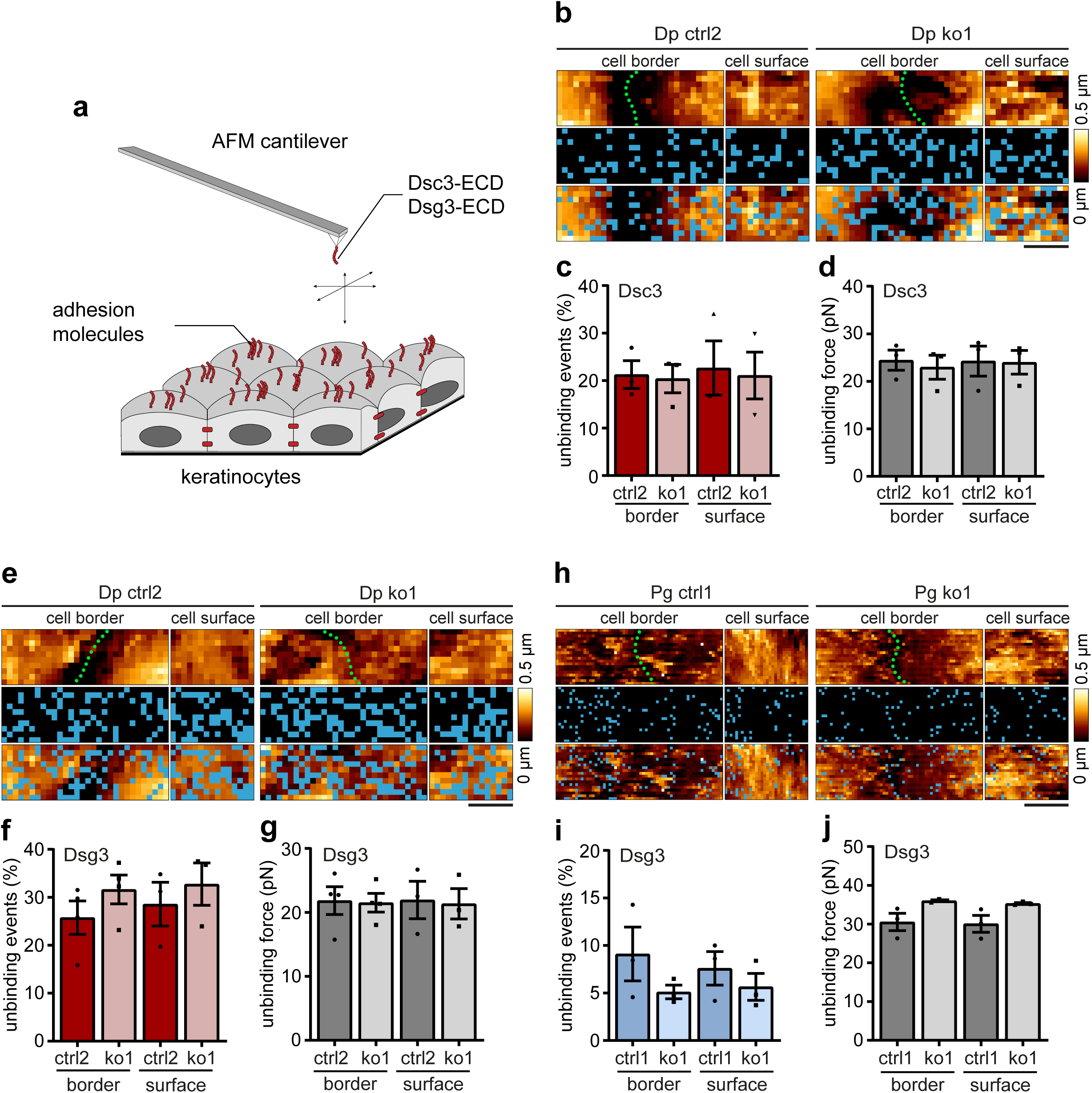
Dsc3 and Dsg3 single molecule binding interactions remain unaffected under loss of Dp and Pg. **(a)** Principle of AFM measurements. A sharp tip on a flexible cantilever was functionalized with recombinant Dsc3 or Dsg3 extracellular domains (ECD) to measure the interaction with molecules on cell surface and cell border areas of living HaCaT keratinocytes. AFM measurements on Dp ctrl and ko cells performed with (**b-d**) Dsc3 or (**e-g**) Dsg3-coated tips to analyze (**b, e**) distribution of unbinding events in adhesion maps, with each blue pixel representing one unbinding event; green dotted lines mark cell borders, (**c, f**) number of unbinding events, and (**d, g**) unbinding force. (**h-j**) AFM measurements on Pg ctrl and ko cells with Dsg3-coated tips to analyze (**h**) distribution of unbinding events in adhesion maps, (**i**) number of unbinding events, (**j**) unbinding force. Each data point represents the mean value of one biological replicate with >225 analyzed force-distance curves, n≥3, error bars represent mean ± SEM. Scale bar: 2 µm.

First, we analyzed the binding properties of Dsc3 due to its profound reduction in Dp ko cells (Fig. 1a). Surprisingly, the number of unbinding events under loss of Dp was not decreased and remained unchanged in comparison to ctrl (Fig. 2c). In addition to that, distribution of single molecule Dsc3 interactions, here depicted by blue pixels in the adhesion map, remained unaltered (Fig. 2b). Importantly, unbinding forces were also unchanged in Dp-deficient cells (Fig. 2d). These experiments were also repeated with a second clone of each cell line, yielding similar results (Supp. Fig. 2). This suggests that Dsc3-dependent interactions are not altered by loss of Dp. In the next setup, we functionalized the AFM tip with Dsg3. Similar to Dsc3, Dsg3-dependent distribution of unbinding events (Fig. 2e) as well as unbinding forces (Fig. 2g) were unaltered, and only a slight increase in the number of unbinding events was detectable in two different Dp ko and ctrl clones (Fig. 2f, Supp. Fig. 2). Because Pg-deficient cells demonstrated reduced levels of Dsg3 (Fig. 1a), we also performed measurements with Dsg3-functionalized tips on Pg ko and corresponding ctrl cells. While the number of unbinding events was only slightly decreased, their distribution remained unchanged in Pg ko (Fig. 2h, i). In addition, unbinding forces were not significantly altered but tended to be increased on both cell border and cell surface areas (Fig. 2j).

In AFM studies, single molecule binding properties are determined by force-distance curves which outline the forces acting on the cantilever as a function of the distance to the sample during retraction of the cantilever. Interaction events are characterized by a jump in force, while the height of the jump represents the unbinding force of the cadherin bond (Fig. 3a). As described previously, the slope preceding the rupture event allows conclusions on the anchorage of the binding partner on the cell surface to the cytoskeleton (Friedrichs et al., 2013). A nonlinear, negative slope indicates a rigid anchorage of the binding partner to the cytoskeletal network (bent unbinding event) due to an interaction event in which the cantilever bends downwards during retraction (Fig. 3a). In contrast, a largely horizontal slope represents the formation of a membrane tether in which the surface protein is pulled up together with the surrounding membrane (plateau unbinding event) (Fig. 3a), suggesting an interaction partner with no anchorage to the cytoskeleton. Measurements with Dsc3-coated cantilevers resulted in similar relative proportions of bent and plateau unbinding events on the cell surface of Dp ctrl and ko clones (Fig. 3b, Supp. Fig. 2). However, we observed a shift from bent to plateau interactions on the cell surface of Dp ko cells using a Dsg3-coated tip (Fig. 3c, Supp. Fig. 2), indicating reduced cytoskeletal anchorage of Dsg3 interaction partners. Together, our data suggest that, in contrast to loss of keratins, the compromised intercellular adhesion in Dp and Pg-deficient cells is not caused by altered interaction probabilities of desmosomal cadherins.

**Fig. 3.**
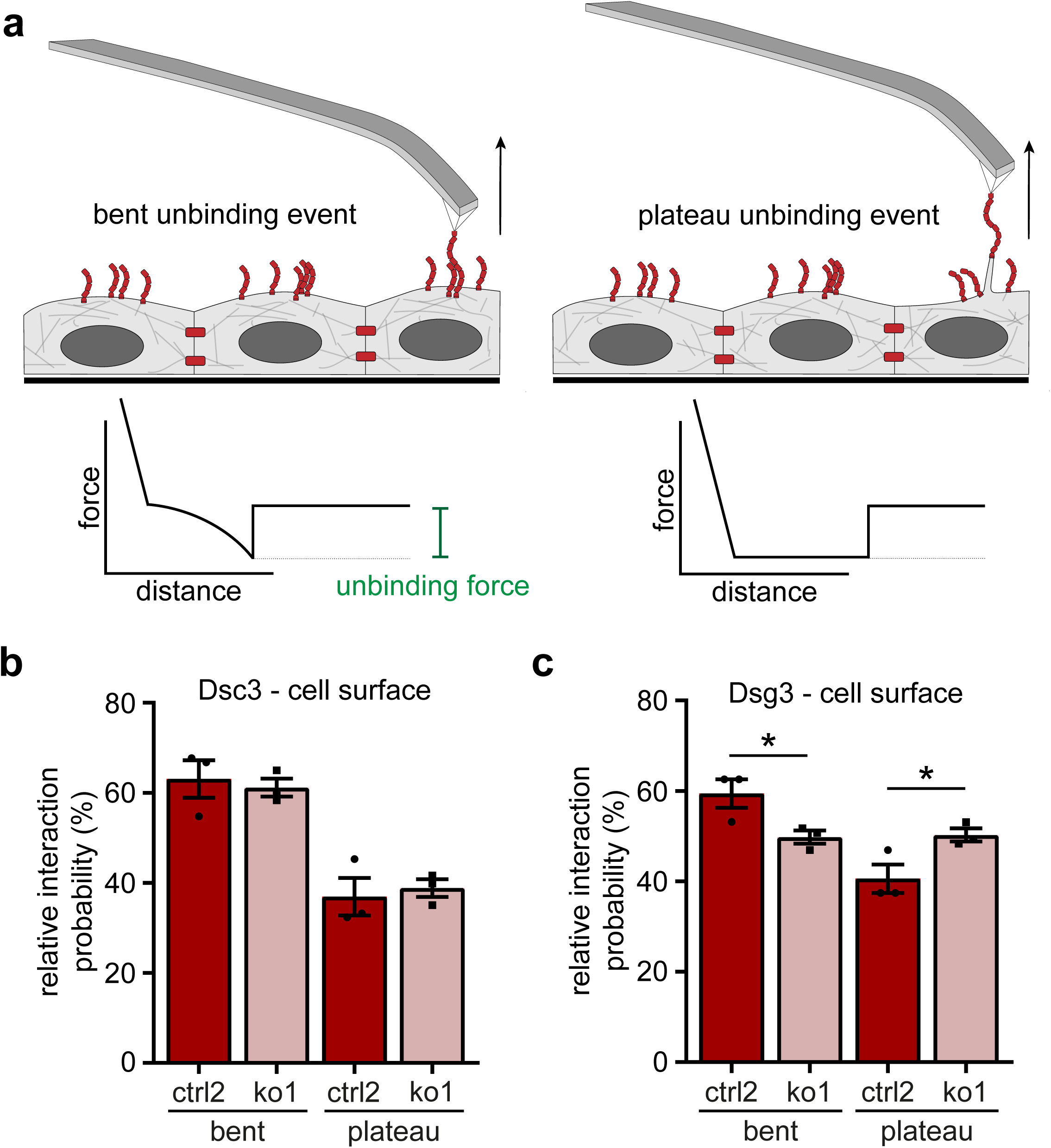
Loss of Dp leads to reduced cytoskeletal anchorage of Dsg3 binding partners. (**a**) Schematic of bent and plateau unbinding events, characterized by appearance of the slope. (**b, c**) AFM measurements with (**b**) Dsc3- or (**c**) Dsg3-coated cantilevers to analyze the relative interaction probability, classified into bent and plateau unbinding events. Each data point represents the mean value of one biological replicate with >225 analyzed force-distance curves, n=3, \**p*<0.05 vs. respective ctrl, error bars represent mean ± SEM.

### Crucial cellular functions remain unaltered under loss of Dp

We further focused on the Dp ko and ctrl cell lines as these gave us the unique opportunity to directly compare human keratinocytes with and without desmosomes. Given this drastic change, we characterized the effects of Dp on crucial cellular functions such as proliferation, apoptosis and cell motility. Using MTT assays, no significant differences in proliferation were observed (Fig. 4a). The percentage of apoptotic cells as determined by TUNEL assays did not reveal significant alterations between Dp ko and ctrl cell lines (Fig. 4b). In previous studies we observed that loss of Dsg2 or Dsg3 regulates cellular migration (Hutz et al., 2017, Rötzer et al., 2016). Thus, cell motility was examined in wound healing assays under basal conditions. Although Dp ko cells migrated slower 8 hours after initial application of the scratch, wound closure accelerated to levels comparable to ctrl lines after 24 hours (Fig. 4c). Taken together, our findings demonstrate a major role of Dp for cell-cell adhesion but revealed no significant contribution to the key cellular functions tested here.

**Fig. 4.**
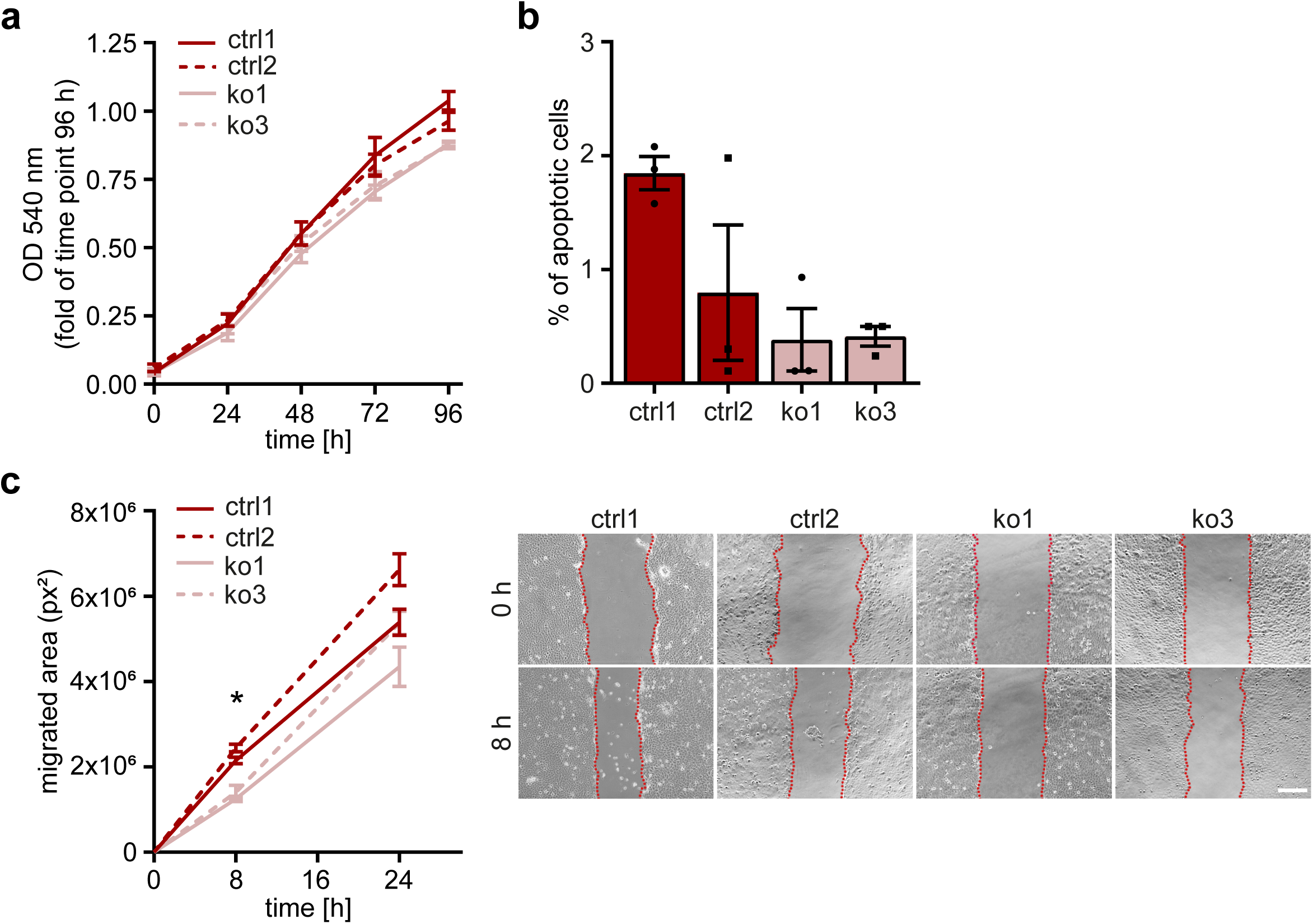
Proliferation, apoptosis and migration remain unaltered under loss of Dp. (**a**) MTT assays in HaCaT Dp ctrl and ko cells. Proliferation was analyzed every 24 h over a time course of 96 h. Optical density of dissolved formazan crystals was measured at 540 nm. n=4, error bars represent mean ± SEM. (**b**) Analysis of apoptotic cells via flow cytometry using TUNEL assay in Dp ko and ctrl cell lines. Each data point represents one biological replicate, n=3, error bars represent mean ± SEM. (**c**) Quantification of scratch wound assays in Dp ko and ctrl cells with representative bright-field images at 0 and 8 h time points. n=5, \**p*<0.05, error bars represent mean ± SEM. Scale bar: 250 µm.

### Dp is required to form large clusters of desmosomal cadherins

With the exception of Dsc3, all other desmosomal cadherins were normally expressed in Dp ko cells. We thus explored whether loss of Dp changed the distribution of desmosomal cadherins at cell-cell interfaces. Immunostainings revealed that Dsc3 and in particular Dsg2 localizes in prominent, punctate clusters at the membrane and co-localizes primarily with Dp in ctrl cells (Fig. 5a). Interestingly, Dp-deficient cell lines displayed a more homogenous distribution of Dsg2 and Dsc3 along the membrane (Fig. 5a). These proteins were subsequently investigated in more detail using structured illumination microscopy (SIM). In Dp ctrl cells, Dsg2 and Dsc3 showed predominantly large, co-localizing clusters with Dsc3 also partially being present outside these clusters. In Dp ko, these larger accumulations were absent and instead appeared to be scattered into discrete small clusters (Fig. 5b). This redistribution of Dsg2 and Dsc3 in the absence of Dp suggests that these molecules mainly occur desmosome-bound and are clustered by Dp at sites of cell-cell adhesion. In contrast, Dsg3 appeared evenly distributed along the membrane in both ko and ctrl cells (Fig. 5a, b). In line with this unaltered distribution, SIM analysis of Dsg3 clusters on the cell surface revealed no differences regarding cluster number and size (Fig. 5c). The correlation between membrane mobility of Dsg3 and the presence of a regularly formed keratin network has been shown previously (Vielmuth et al., 2018). Nevertheless, fluorescence recovery after photo-bleaching (FRAP) at the interfaces of two adjacent Dsg3-GFP expressing cells revealed similar mobility of Dsg3 in Dp ctrl and ko cells (Fig. 5d). This indicates that Dsg3 membrane turnover is also not affected by loss of Dp. These experiments revealed that different desmosomal cadherins have distinct distributions within the membrane, with Dsg2 and to lesser extents Dsc3 being largely restricted to desmosomes and Dsg3 being distributed both inside and outside of desmosomes. Furthermore, Dp is required to form large clusters of cadherins.

**Fig. 5.**
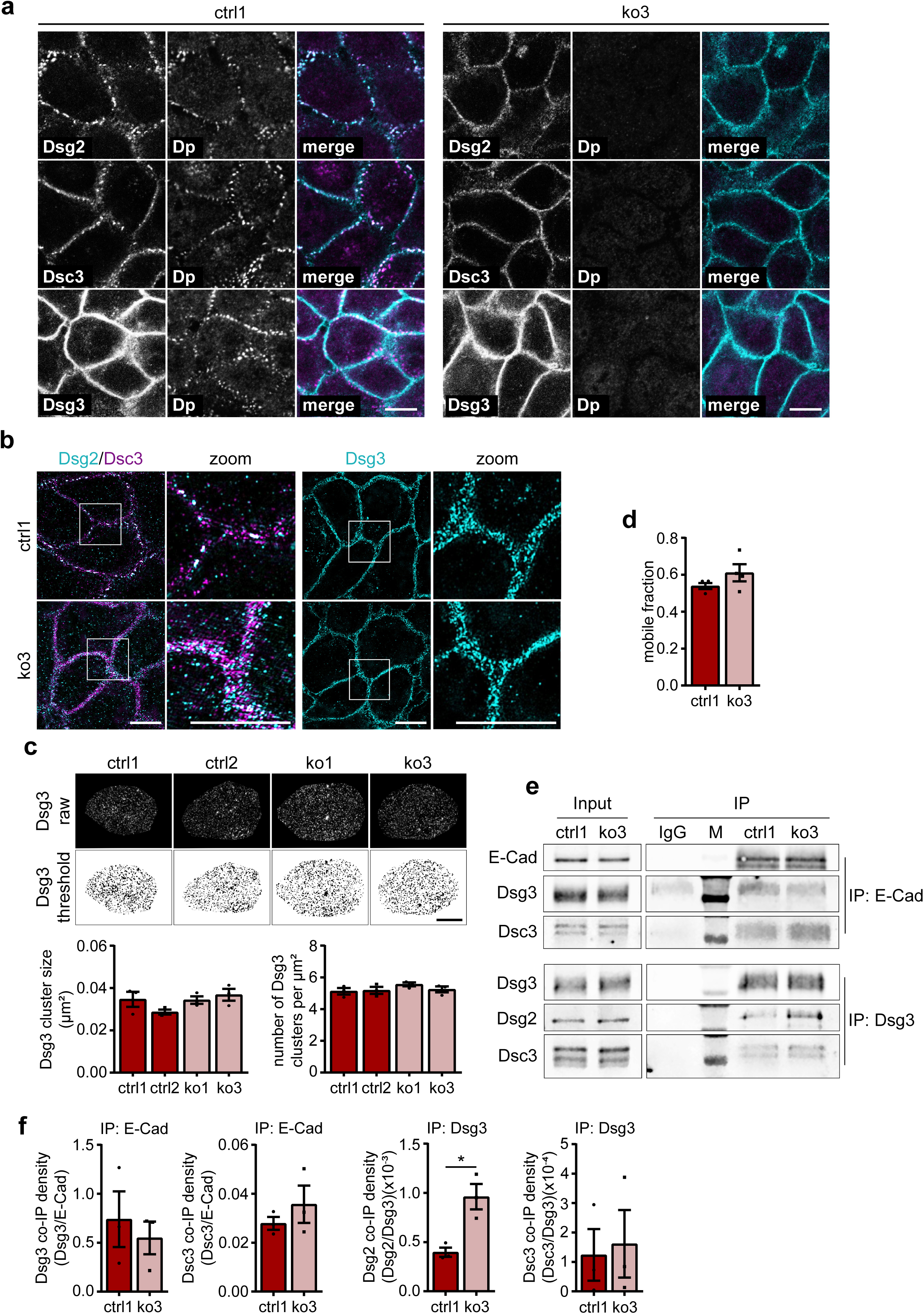
Redistribution of Dsg2 and Dsc3, but not Dsg3 occurs under loss of Dp. (**a**) Representative immunostainings of Dp (magenta), double-stained with either Dsg2 or Dsc3 (turquoise) in Dp ctrl (left panel) and ko (right panel) cells, n=3. Scale bar: 10 µm. (**b**) SIM images to visualize Dsg2, Dsc3 and Dsg3 in Dp ctrl and ko cells. White boxes mark the magnified areas, n=3. Scale bar: 10 µm. (**c**) SIM images of apical cell portions in Dp ctrl and ko cells. Background was subtracted and a threshold was used to display Dsg3 clusters (upper panel). Quantification of Dsg3 cluster size and cluster number (lower panel) obtained from SIM images. n=3, error bars represent mean ± SEM. Scale bar: 5 µm. (**d**) Quantification of FRAP Dsg3-GFP experiments in Dp ctrl and ko cells. Each data point represents the mean value of the mobile fraction of Dsg3-GFP of 2-8 bleached cell borders of one independent experiment, n=4, error bars represent mean ± SEM. (**e**) Immunoprecipitations (IPs) of E-Cad and Dsg3 in Dp ctrl and ko cells, M=marker, n=3. IP with species matched IgG served as control. (**f**) Quantification of co-IPs from E-Cad and Dsg3-IP shown in (**e**); band densities were normalized to the respective IP band density, n=3, \**p*<0.05 vs. ctrl, error bars represent mean ± SEM.

The first steps of desmosome assembly were shown to require binding of E-Cad to Dsgs (Rotzer et al., 2015, Shafraz et al., 2018). As desmosomal cadherins were still located at the membrane in Dp ko but no desmosomes were formed, we evaluated whether these complexes were affected in Dp-deficient cells. Co-immunoprecipitations revealed interactions of E-Cad with Dsg3 and Dsc3 to similar extents in Dp ko as well as in ctrl cell lines (Fig. 5e, f), indicating regular formation of desmosomal precursors under loss of Dp. Interestingly, we did not detect interactions of E-Cad with Dsg2 using two different antibodies (data not shown). By immunoprecipitating Dsg3, we found increased interactions of Dsg3 with Dsg2, but not Dsc3, in Dp ko cells (Fig. 5e, f), suggesting that normally desmosome-restricted Dsg2 molecules cluster with Dsg3 when desmosomes are not present.

Our results indicate that loss of Dp did not interfere with the formation of early desmosomal precursors and membrane availability of desmosomal cadherins. Instead, redistribution of Dsg2 and Dsc3 along the membrane of cells lacking desmosomes demonstrates the role of Dp in forming large clusters of desmosomal cadherins to maintain intercellular adhesion. To examine whether clustering requires Dp expression in both adjacent cells, we seeded mixed populations of Dp ctrl and ko cells and allowed monolayer formation. Dp ctrl cells were labeled with CellTrace Violet (CTV) to allow a distinction from unlabeled Dp ko cells. SIM revealed the formation of large sandwich-like clusters between ctrl cells, consisting of Dsg2 flanked by Dp in both cells (Fig. 6a orange boxes). Interestingly, Dp and Dsg2 also formed large clusters at the interface of ctrl and ko cells, yielding a split-desmosome like appearance (Fig. 6a green boxes). Albeit not being present at all ctrl-ko interfaces, the size and appearance of these Dsg2 clusters seemed to be largely similar to those clusters found at ctrl-ctrl interfaces. These findings suggest that Dp expression in one of two adjacent cells is sufficient to cluster Dsg2 and that Dp is able to accumulate at sites of cell-cell contact, independent of Dp expression in the neighboring cell.

**Fig. 6.**
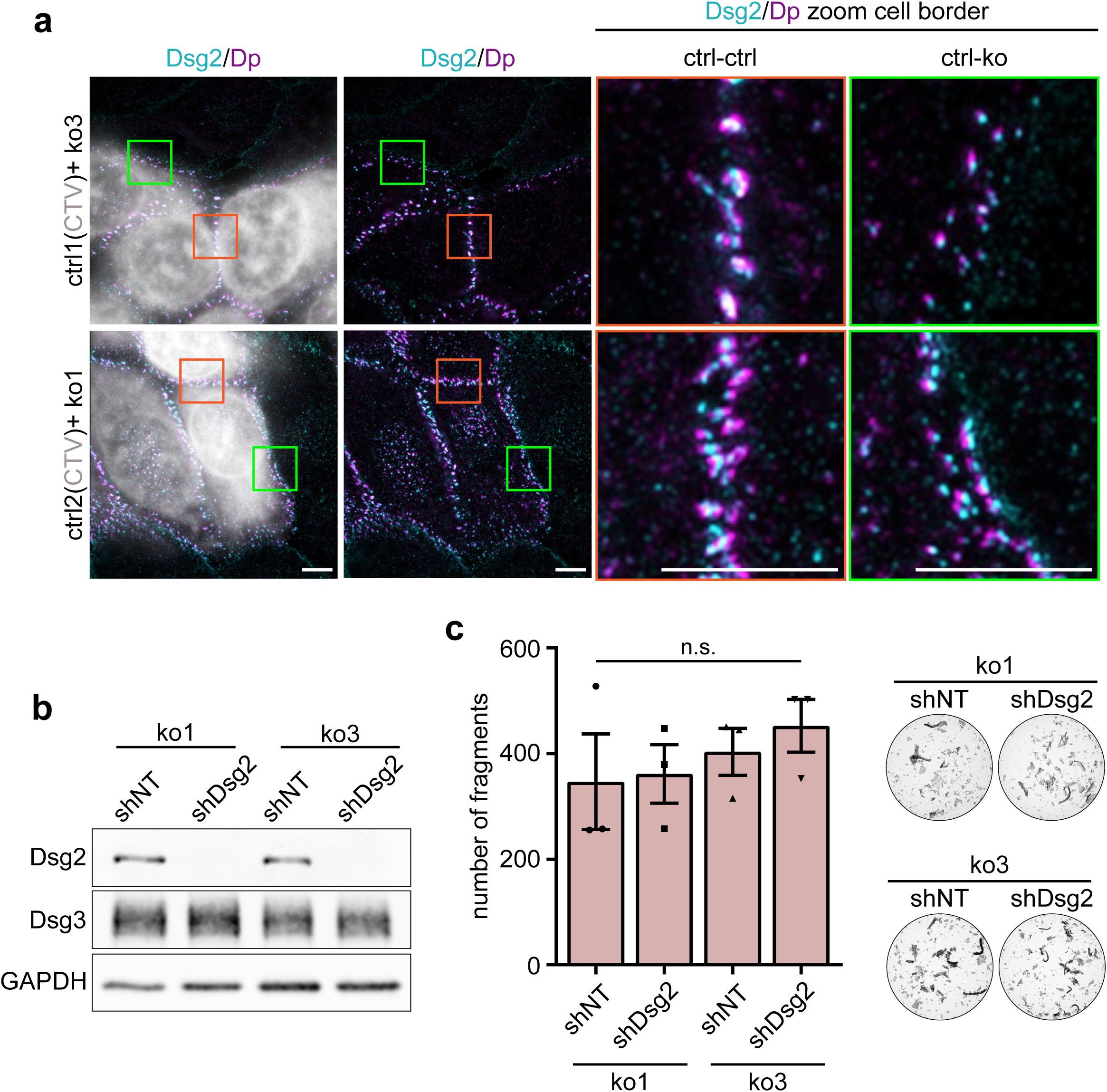
Downregulation of desmosomal cadherins in Dp ko does not further decrease intercellular adhesion. (**a**) Maximum intensity projections acquired by SIM. Dp ctrl cell lines were stained with CTV (shown in gray) as marker and cultured together with unstained Dp ko cells. Mixed monolayers were subsequently stained for Dsg2 and Dp. Orange boxes mark the magnified areas containing ctrl-ctrl cell borders. Green boxes mark the magnified areas containing ctrl-ko cell borders. n=3. Scale bar: 5 µm. (**b**) Confirmation of shDsg2-mediated knockdown in Dp ko cell lines via Western blot analysis. shNT-transduced cells were used as controls. GAPDH served as a loading control, n=3. (**c**) Representative images and quantification of Dp ko shNT and shDsg2 monolayers after dissociation assay. Each data point represents the mean value of ≥2 replicates of one independent experiment, n=3, \**p*<0.05, error bars represent mean ± SEM.

### Silencing of desmosomal cadherins does not further decrease intercellular adhesion in Dp ko

To finally investigate whether the small clusters of desmosomal cadherins in Dp ko contribute to intercellular adhesion, Dsg2 was silenced in Dp ko cell lines by shRNA. Although strong downregulation of Dsg2 was achieved (Fig. 6b), intercellular adhesion in Dp ko was not further decreased in response to shear stress (Fig. 6c). This suggests that extradesmosomal cadherin clusters do not significantly contribute to cell-cell adhesion but need to be clustered within desmosomes by Dp to facilitate strong cell-cell adhesion.

## DISCUSSION

### Dp is indispensable for desmosome formation and intercellular adhesion

We here show that loss of Pg in human keratinocytes resulted in morphologically altered desmosomes and weakened intercellular adhesion. In contrast, Dp-deficient cells failed to form desmosomes, resulting in a massive loss of cell-cell adhesion. This demonstrates that Dp is required to form desmosomes and underscores the relevance of these complexes for intercellular adhesion. Previous studies showed that cultured keratinocytes derived from a Pg ko mouse model displayed less desmosomes with exhibition of compromised cytoplasmic plaques (Acehan et al., 2008), while global deletion of Dp had a more drastic impact and led to severely decreased desmosome numbers and embryonic death before E5 (Gallicano et al., 1998). Although an epidermis-specific knockout model still displayed desmosome numbers comparable to wildtype (wt) mice, the animals died shortly after birth due to sloughing of the epidermis (Vasioukhin et al., 2001), which demonstrates severely impaired desmosomal functions. One common theme in mouse models with deletions of Dp or Pg appears to be the reduced attachment of intermediate filaments (Acehan et al., 2008, Gallicano et al., 1998, Li et al., 2012, Vasioukhin et al., 2001). This suggests that the connection to intermediate filaments is crucial for intercellular adhesion and it is well established that anchorage to the desmosomal plaque is dependent on both Pg and Dp (Acehan et al., 2008, Bornslaeger et al., 1996). Still, the phenotype in Dp ko models is worse compared to those models lacking other plaque proteins, which was also reproduced in our Dp and Pg ko cell lines. This indicates that additional mechanisms fail in response to Dp loss. While other studies implicate Dp in the regulation of proliferation and apoptosis (Wan et al., 2007, Yang et al., 2012), which may contribute to downregulation of junctions, we did not observe consistent changes in our models. Additionally, cellular migration was not altered in Dp ko cells, while it was increased in epidermal cancer cells upon downregulation of Dp (Bendrick et al., 2019). This indicates that these changes are context-dependent but do not contribute to the massive loss of cell adhesion in response to Dp depletion. We thus tested several other possibilities how Dp may modulate desmosome formation and intercellular adhesion in addition to compromised keratin anchorage. These possibilities included changes (i) in the binding properties of individual desmosomal cadherins (Vielmuth et al., 2018), (ii) in the formation of nascent desmosomes and (iii) in the clustering of desmosomal cadherins on the cell surface as further discussed below.

Cell adhesion is a highly plastic process and compensatory effects in response to depletion are possible. For instance, loss of one desmosomal cadherin can be ameliorated by endogenous upregulation of another desmosomal cadherin (Hartlieb et al., 2014, Walter et al., 2019). Loss of Pg can result in upregulation and partial compensation by β-Cat which was also the case (albeit not significant) in our Pg ko cells. In contrast, no compensatory upregulation of any other desmosomal molecule was detectable in Dp ko cells, which may additionally contribute to the more pronounced loss of cell adhesion in Dp ko compared to Pg ko cells.

### Dp does not regulate interaction properties but is required for large cluster formation of desmosomal cadherins

Our data demonstrate that interaction forces of individual desmosomal cadherins were neither altered by loss of Dp nor by loss of Pg. This is surprising as in cells lacking all keratins, the interaction forces of single Dsg3 molecules were reduced (Vielmuth et al., 2018), suggesting that the linkage to the intermediate filament cytoskeleton regulates desmosomal cadherin binding properties. However, it was also shown that these changes required the activity of signaling molecules. Thus, it appears possible that the mechanism is indirect and not dependent on direct interactions between keratins and the desmosomal plaque. In support, it was shown that increased phosphorylation of Pg increases forces of Dsg2-dependent interactions in cardiac myocytes, indicating that kinase-dependent modifications rather than cytoskeletal interactions modulate individual binding properties (Schinner et al., 2017). Because we saw a scattering of remaining Dsc3 and Dsg2 molecules over the entire membrane of Dp ko cells, which were largely restricted to desmosomes in ctrl cells, it is likely that these free molecules now contribute to the binding events detected by AFM. In support, Dsc3 binding frequency remained unaltered and we even detected a slight increase of Dsg3 single molecule interactions in Dp ko. This suggests that both homophilic as well as heterophilic (e.g. Dsg3-Dsg2 or Dsc3-Dsg2) interactions take place on the free surface of keratinocytes. In line with this, previous studies suggest not only homo-but, heterophilic interactions between Dsg and Dsc isoforms (Harrison et al., 2016, Spindler et al., 2009), specifically between Dsg2 and Dsg3 (Vielmuth et al., 2018).

Together, these data do not support a concept in which the massive loss of intercellular adhesion can be explained by differences in the binding properties of individual desmosomal molecules. De-novo formation of desmosomes can be dissected into discrete steps (Moch et al., 2020, Nekrasova and Green, 2013). Desmosomal cadherins are transported to the membrane independent of Dp (Nekrasova et al., 2011). At the membrane, clustering with E-Cad appears to be required, followed by segregation into desmosomal complexes later on (Rotzer et al., 2015, Shafraz et al., 2018). Our immunoprecipitation data demonstrate that E-Cad interacts with Dsg3 and Dsc3 to similar extents in ctrl and ko cell lines, indicative of regular initial steps of desmosome formation. Live cell imaging studies suggest that Dp from cytosolic pools subsequently locates to nascent desmosomal cadherin clusters (Godsel et al., 2005, Moch et al., 2020). In view of the observation that the Dp ko cells generated in this study do not exhibit ultrastructurally detectable desmosomes but rather show desmosomal cadherins scattered in small clusters in the membrane, it appears that Dp is required to coalesce and stabilize these small clusters into larger assemblies, finally yielding mature desmosomes. Importantly, depletion of Dsg2 in Dp ko cells did not further destabilize intercellular adhesion. This demonstrates that extradesmosomal cadherins do not significantly contribute to cell cohesion and fusion of small clusters into desmosomes is required in this regard. As desmosomes were shown to organize the keratin network adjacent to the plasma membrane (Moch et al., 2020), it is possible that Dp is an important organizing center for both desmosomal cadherins and keratins. Accordingly, in contrast to Dp, cells lacking all keratin filaments still form morphologically compromised desmosomes (Kroger et al., 2013). Whether Dp is required to segregate intermediate complexes containing both classical and desmosomal cadherins during desmosome assembly remains to be further addressed. Accumulation of these complexes in Dp ko cells was not detectable, which may be either due to a continuous degradation or due to normal segregation with subsequent failure of large cluster formation. Interestingly, large clustering and correct positioning of Dp can take place even if the adjacent cell lacks Dp. This suggests that trans-interaction of desmosomal cadherins is a critical step in desmosome formation. This is in line with a previous study showing that microprinting of Dsc2 was sufficient to recruit Dp to the membrane in these areas (Lowndes et al., 2014).

Taken together, classical and desmosomal cadherins, desmosomal plaque proteins as well as keratin filaments all appear to contribute in multiple spatiotemporally distinct as well as interdependent steps to desmosome formation. Dp appears to be crucial in this regard, and large-scale clustering of desmosomal cadherins dependent on Dp appears to be critical for both desmosome formation and intercellular adhesion.

## MATERIALS AND METHODS

Please refer to the supplementary material.

## ABBREVIATIONS

AFM: atomic force microscopy;
ctrl: control;
CTV: CellTrace Violet;
Dp: desmoplakin;
Dsc: desmocollin;
Dsg: desmoglein;
E-Cad: E-cadherin;
FRAP: fluorescence recovery after photo bleaching;
ko: knockout;
Pg: plakoglobin;
Pkp: plakophilin;
SIM: structured illumination microscopy;
wt: wildtype;
β-Cat: β-catenin;

## CONFLICT OF INTEREST

The authors state no conflict of interest.

## ACKNOWLEDGEMENTS

We are grateful to Carola Alampi and Dr. Cinzia Tiberi from the Center for Cellular Imaging and NanoAnalytics (C-CINA, University of Basel, Switzerland) and Dr. Alexia Loynton-Ferrand from the Imaging Core Facility (IMCF, University of Basel, Switzerland) for best support. We thank Dr. Mariya Radeva (Faculty of Medicine, Ludwig-Maximilians-Universität Munich, Germany) for helpful scientific discussions as well as Anja Fuchs, Martina Hitzenbichler, Sabine Mühlsimer, Kilian Skowranek and Aude Zimmermann for excellent technical assistance. This project was supported by German Research Council projects SP1300-1/3 and SP1300-3/1.

