## Supplementary Material for "Clustering of desmosomal cadherins by desmoplakin is essential for cell-cell adhesion"

### MATERIALS AND METHODS

#### Cell Culture and Generation of CRISPR/Cas9 Cell Lines

Spontaneously immortalized HaCaT (Human adult high Calcium low Temperature) keratinocytes (Boukamp et al., 1988) were cultured in a humidified atmosphere of 5% $CO_2$ and 37°C in Dulbecco's Modified Eagle Medium (DMEM) (Sigma-Aldrich, St. Louis, MO, USA) containing 1.8 mM $Ca^{2+}$ and complemented with 10% fetal bovine serum (Merck, Darmstadt, Germany), 50 U/ml penicillin, 50 µg/ml streptomycin (both AppliChem, Darmstadt, Germany) and 4 mM L-glutamine (Sigma-Aldrich). Pg and Dp ko and ctrl cell lines were generated using CRISPR/Cas9. Each protein was targeted separately in two different exons of the respective human gene using plasmids containing the expression vector pCMV-Cas9-GFP (purchased from Sigma-Aldrich). The following target sites were chosen: *JUP* exon 2 (CTTGATAGGCTGCTCC ATCAGG) or exon 3 (AGGGCCAAACGGGTGCGGGAGG), *DSP* exon 1 (AGAGTGTTGA TCCGCGGGTGGG) or exon 3 (TTGGATGAGTGTTTTGCCCAGG). HaCaT cells were seeded in 6-well plates and transiently transfected the following day at 70-90% confluency using Lipofectamine 2000 (Thermo Fisher Scientific, Waltham, MA, USA) according to the manufacturer's protocol. After 24 h, cells expressing GFP (thus demonstrating the presence of Cas9) were single cell sorted into 96-well plates by a FACSAria III (BD Biosciences, Franklin Lakes, NJ, USA) cell sorter. To promote cellular growth, single cell colonies were cultured with sterile filtered, 50% conditioned medium collected from confluent HaCaT wildtype (wt) cell culture flasks. Expanded clones were validated by Western blot, immunostaining and sequencing of genomic DNA. For each protein, three ko and three ctrl cell lines were chosen for experiments.

#### Generation of Lentiviral Constructs and Stable Cell Lines

A Small hairpin (sh) Dsg2 knockdown (kd) construct was purchased from Merck Millipore (MISSION shRNA construct, shDsg2: #TRCN53843, seq.: CCGGGCTCAAACT AACGAAGGAATTCTCGAGAATTCCTTCGTTAGTTTGAGCTTTTG). A construct with a non-target sequence (a gift from David Sabatini, Addgene plasmid #1864; http://n2t.net/addgene:1864; RRID: Addgene_1864, seq.: CCTAAGGTTAAGTCGCCCT CGCTCGAGCGAGGGCGACTTAACCTTAGG) was used as control. For lentivirus generation, HEK293T cells were co-transfected overnight with the packaging vector psPAX2, the envelope vector pMD2.G (both a gift from Didier Trono, Addgene plasmid #12259; http://n2t.net/addgene:12259; RRID: Addgene_12259 and Addgene plasmid #12260; http://n2t.net/addgene:12260; RRID: Addgene_12260) and kd construct using TurboFect (Thermo Fisher Scientific). Medium was changed and supernatant containing virus particles was collected and enriched using LentiConcentrator (OriGene, Rockville, MD, USA) after 48 h. Transduction of HaCaT cell lines was carried out using 5 µg/ml Polybrene (Sigma-Aldrich) according to the manufacturer's instructions. After 24 h, medium was changed and 2 µg/ml Puromycin (Thermo Fisher Scientific) was used 48 h post-transduction for selection and maintenance of clones. Knockdown was validated via Western blot.

#### Western Blot

Confluent HaCaT cells in 24-well plates were washed with phosphate-buffered saline (PBS) and scraped in SDS lysis buffer (25 mM HEPES, 2 mM EDTA, 25 mM NaF, 1% SDS, pH 7.6) supplemented with an equal volume of a protease inhibitor cocktail (cOmplete, Roche Diagnostics, Mannheim, Germany). Lysates were sonicated and the total protein amount was determined with a BCA protein assay kit (Thermo Fisher Scientific) according to the manufacturer's instructions. After lysates were denaturized for 5 min at 95°C in Laemmli buffer, gel electrophoresis and wet blotting on nitrocellulose membranes (Thermo Fisher Scientific) were carried out according to standard procedures. Membranes were blocked in Odyssey blocking buffer (Li-Cor, Lincoln, NE, USA) for 1 h at room temperature. The following primary antibodies were diluted with 5% BSA in tris-buffered-saline containing 0.1% Tween20 (TBS-T) (Thermo Fisher Scientific) and applied overnight at 4°C: mouse anti-Dsc2 mAb (#60239-1-Ig Proteintech, Rosemont, IL, USA), mouse anti-Dsc3 mAb (clone U114, #65193, Progen, Heidelberg, Germany), mouse anti-Dsg2 mAb (clone 10G11, #BM5016 Acris, Herford, Germany), rabbit anti-Dsg3 pAb (EAP3816 Elabscience, Biozol, Eching, Germany), mouse anti-Pkp1 mAb (clone 10B2, #sc-33636 Santa Cruz, Dallas, TX, USA), mouse anti-Pkp2 mAb (#651101 Progen), mouse anti-Pkp3 mAb (#651113 Progen), mouse anti-Pg mAb (clone PG5.1, #61005 Progen), mouse anti-Dp mAb (#sc-390975 Santa Cruz), mouse anti-E-Cad mAb (clone 36, #610181 BD Biosciences), mouse anti-β-Cat mAb (clone 14, #610154 BD Biosciences), mouse anti-GAPDH mAb (clone 0411, #sc-47724 Santa Cruz). As secondary antibodies, goat anti-mouse 800CW and goat anti-rabbit 680RD (#925-32210 and #925-68071, both Li-Cor) were incubated for 1 h at room temperature. Protein bands were detected in an Odyssey FC imaging system and band density was quantified with ImageStudio (both Li-Cor).

**Immunoprecipitation**

Cells were grown to confluency, washed twice with ice-cold PBS and incubated for 30 min on ice with modified RIPA buffer (10 mM $Na_2HPO_4$, 150 mM NaCl, 1%Triton X-100, 0.25% SDS, 1% sodium deoxycholate, pH 7.3) complemented with aprotinin, leupeptin, pepstatin and phenyl-methylsulphonyl fluoride. Subsequently, lysates were scraped and homogenized on ice by passing ten times through a 20G and afterwards a 25G injection needle. Samples were centrifuged (7.700x *g*, 5 min, 4°C) and protein concentration in the supernatant was determined using the BCA method (Thermo Fisher Scientific). Equal amounts of protein were incubated with the following antibodies overnight at 4°C: mouse anti-Dsg2 mAb (clone 10G11, #BM5016 Acris), mouse anti-Dsg1/2 mAb (clone DG 3.10, #61002 Progen), rabbit anti-Dsg3 pAb (EAP3816 Elabscience), mouse anti-Dsc3 mAb (clone U114, #65193, Progen), mouse anti-E-Cad mAb (clone 36, #610181 BD Biosciences), normal mouse IgG (#sc-2025) and normal rabbit IgG (#sc-2027, both Santa Cruz). Lysates were incubated with pre-washed, magnetic Dynabeads Protein G (Thermo Fisher Scientific) for 1 h at 4°C. Beads were washed twelve times and afterwards mixed with Laemmli buffer and denaturized at 95°C for 10 min. Eventually, the whole sample was loaded on SDS polyacrylamide gels and electrophoresis, wet blotting and protein band detection was carried out as described in the section *Western Blot*.

**<u>Cell Dissociation Assay</u>**

24 h after reaching confluency, HaCaT keratinocytes were washed with pre-warmed HBSS and incubated with 150 µl dispase II (>2.4 U/ml in HBSS; Sigma-Aldrich) for 20 min at 37°C to detach the intact cell sheet from the well bottom. Subsequently, the dispase solution was replaced by 350 µl HBSS containing 0.5 mg/ml thiazolyl blue tetrazolium bromide (MTT; VWR, Radnor, PA, USA) to improve visibility of the monolayer. Mechanical shear stress was applied three times using an electrical pipette (Eppendorf, Hamburg, Germany). Wells were finally imaged with a binocular microscope (Olympus, Tokyo, Japan) and SLR camera (Canon, Tokyo, Japan). Cell fragments, which are a direct measure for impaired cell cohesion, were counted manually using ImageJ (National Institutes of Health, Bethesda, USA).

**Electron Microscopy**

Cells were seeded on 24 mm glass coverslips. After reaching confluency, cells were fixed by adding an equal volume of pre-warmed 5% glutaraldehyde (GA) + 4% paraformaldehyde (PFA) (Electron Microscopy Sciences, Hatfield, PA, USA) in 0.1 M Pipes buffer with 2 mM $CaCl_2$ (pH 7.3) to the culture medium for 15 min at room temperature. The solution was replaced by 2.5% GA + 2% PFA with 2 mM $CaCl_2$ in 0.1 M Pipes buffer for 2 h at room temperature and subsequently incubated for 16 h at 4°C. After fixation, cells were washed three times with cold 0.1 M Pipes with 2 mM $CaCl_2$ and then rinsed several times with 0.1 M cacodylate buffer (pH 7.3). Cells were post-fixed for 1 h at 4°C using 1% osmium tetroxide + 0.8% potassium ferracyanide (Electron Microscopy Sciences) in 0.1 M cacodylate buffer. After several washes with cacodylate buffer and ultrapure distilled water, coverslips were *en bloc* stained in the dark with 1% aqueous uranyl acetate (Electron Microscopy Sciences) for 1 h at 4°C. For dehydration of the cells, an ascending ethanol series was used at 4°C. After three washes with absolute ethanol, cells were rinsed in acetone and first embedded in a mixture of resin/acetone followed by pure Epon 812 resin (Electron Microscopy Sciences) overnight. Cell-side down, samples were mounted on BEEM capsules (Electron Microscopy Sciences) filled with EPON. After polymerization at 60°C for 48 h, samples were removed from the EPON block with the nitrogen hot water method. 70 nm thin serial sections, cut with a diamond knife, were mounted on formvar-carbon coated copper slot grids, stained with uranyl acetate and Reynolds`s lead citrate. Samples were examined in a FEI Tecnai T12 spirit Transmission Electron Microscope (Thermo Fisher Scientific) operating at 80 kV equipped with a CCD Veleta digital camera.

#### Immunostaining

Cells were grown on 13 mm glass coverslips and fixed with either 2% PFA (Thermo Fisher Scientific) in PBS at room temperature or ice-cold methanol (Merck Millipore) for 10 min on ice. Cells were permeabilized with 0.1% Triton X-100 in PBS for 5 min and afterwards blocked with 3% BSA and 1% normal goat serum in PBS for 1 h. The following primary antibodies were incubated overnight at 4°C: mouse anti-Dsc3 mAb (clone U114, #65193 Progen), mouse anti-Dsg2 mAb (clone 10G11, #BM5016 Acris), rabbit anti-Dsg2 pAb (#610121 Progen), mouse anti-Dsg3 mAb (clone 5G11, # 326300 Invitrogen, Carlsbad, CA, USA), rabbit anti-Dp pAb (NW6), mouse anti-Dp mAb (1G4) (both kind gifts from Kathleen Green, Northwestern University, Chicago, USA), mouse anti-Pg mAb (clone PG5.1, #61005, Progen). After several washing steps with PBS, cy3 or cy5 conjugated anti-rabbit or anti-mouse antibodies (Dianova, Hamburg, Germany) were incubated for 1 h at room temperature and DAPI (Sigma-Aldrich) was added for 10 min to counterstain nuclei. Samples were mounted with ProLong Diamond Antifade (Thermo Fisher Scientific) and for image acquisition a 63x PL APO NA=1.4 objective on a LSM710 confocal microscope (Zeiss, Oberkochen, Germany) was used.

#### Structured Illumination Microscopy (SIM)

For SIM, immunofluorescence was carried out as described in the section *Immunostaining* with the following modifications: after fixation with ice-cold methanol or 4% PFA for 20 min, cells were permeabilized with 0.5% Triton X-100 in TBS. PBS was always substituted with TBS and 0.1% Triton X-100 in TBS. For experiments with simultaneous seeding of Dp ctrl and ko cells, Dp ctrl cells were labeled with CellTrace Violet (CTV) (Thermo Fisher Scientific). Confluent Dp ctrl monolayers were detached by trypsinization, washed with PBS (200x *g*, 3 min) and incubated with

CTV according to the manufacturer's instructions in a water bath at 37°C for 12 min. Cells were subsequently washed twice with medium, mixed with unlabeled Dp ko cells and seeded on 13 mm glass coverslips. Fixation, blocking and staining were carried out as described above. Images were acquired using a DeltaVision OMX-Blaze (Version 4; Applied Precision, Issaquah, WA) equipped with a 60x PL APO NA=1.42 objective (Olympus) and analyzed with Image J (NIH, Bethesda, MD, USA).

#### Atomic Force Microscopy (AFM)

AFM measurements were performed with a Nanowizard IV AFM (JPK Instruments, Berlin, Germany) mounted on an inverted optical microscope (IX83, Olympus). $Si_3N_4$ MLCT cantilevers (Bruker, Mannheim, Germany) were functionalized with either recombinant Dsg3-ECD or Dsc3-ECD constructs using a bifunctional polyethylene glycol linker (acetal-PEG-NHS, Gruber Lab, Institute of Biophysics, Linz, Austria) as described before (Ebner et al., 2007). Living HaCaT keratinocytes were probed with the pyramidal-shaped D-Tip in standard cell culture medium containing 1.8 mM $Ca^{2+}$ at 37°C. After image acquisition of topographical overviews, regions of interest (ROI) comprising cell surface areas of 3 µm x 3 µm and areas of 7.5 µm x 2.5 µm spanning the borders of two adjacent cells were selected. In ROI, the force mapping mode was used to detect single interactions between molecules immobilized on the tip of the cantilever and molecules on the surface of the cells. Briefly, the functionalized tip was repetitively brought into contact with the cell monolayer at a speed of 10 µm/s. After reaching a defined setpoint of 0.2 nN and a contact time of 0.1 s, the cantilever was retracted again with a Z-length of 2 µm, resulting in one force-distance curve at each predefined pixel. In case of interacting molecules, the cantilever bends downwards during retraction until the bond eventually ruptures and the cantilever jumps back into the neutral position (unbinding event). Analysis of force-distance curves was carried out using the JPKSPM Data Processing software (Version 6, JPK Instruments, Berlin, Germany).

#### MTT Assay

For proliferation assays, $1x10^5$ HaCaT cells were seeded into 24-well plates (0 h). At respective time points (24, 48, 72, 96 h) medium was replaced by fresh medium containing 0.5 mg/ml MTT. Cells were treated for 90 min at 37°C to allow reduction of MTT to purple formazan in viable cells. After two washing steps with PBS, 150 µl dimethyl sulfoxide (DMSO) + 50 µl PBS were added and incubated until complete solubilization of formazan crystals. Supernatants were transferred to a 96-well plate and absorbance was quantified at 540 nm by a microplate reader (Synergy H1, BioTek, Winooski, VT, USA). Wells without cells undergoing the same protocol served as blanks and respective optical density (OD) was subtracted from sample values.

#### Cell Migration Assay

HaCaT clones were seeded in duplicate and grown to confluency in 24-well plates. A sterile pipette tip was used to manually induce one vertical scratch wound per well. After washing with cell culture medium to remove debris, cells were incubated at 37°C for 24 h. To monitor cell motility, scratch wounds were imaged at 0, 8 and 24 h using a bright-field microscope (CKX53 inverted microscope, Olympus). The ImageJ software free hand selection tool (NIH) was used to assess the uncovered scratch area and the migrated area was calculated by subtracting the wound area at respective time points from the initial wound area.

**Terminal Deoxynucleotidyl Transferase-mediated dUTP Nick-End Labeling (TUNEL) Assay**

The DeadEnd Fluorometric TUNEL System (Promega, Madison, WI, USA) was used according to the manufacturer's instructions. Briefly, cells were grown to confluency, detached by trypsinization and washed two times with PBS (300 x *g*, 5 min, 4°C). Cells were fixed with 2% PFA for 20 min on ice. After two additional washing steps with PBS, cells were eventually permeabilized with ice-cold 70% ethanol at -20°C overnight. Next, cells were washed twice in PBS (500 x *g*, 10 min). To obtain positive controls, cells were incubated for 10 min with 100 µl DNase I buffer (10 U/ml; AppliChem) and subsequently washed three times with deionized water. Afterwards, $2x10^6$ cells of each condition were treated with 80 µl equilibration buffer for 5 min at room temperature. Cells were then incubated with 50 µl nucleotide-enzyme mix in a water bath for 60 min at 37°C and the reaction was finally terminated with 20 mM EDTA. After several washing steps with PBS, DAPI was added for 30 min at room temperature to stain nuclei. Analysis of fluorescence was carried out immediately by flow cytometry using a Cytoflex (Beckman Coulter, Brea, CA, USA).

**Fluorescence Recovery After Photobleaching (FRAP)**

HaCaT cells were seeded in 8-well imaging chambers (Ibidi, Martinsried, Germany) and transiently transfected the following day with pEGFP-N1-Dsg3 (kind gift from Yasushi Hanakawa, Ehime University School of Medicine, Japan) using TurboFect. FRAP studies were performed 24 h post-transfection using a LSM710 confocal microscope equipped with a 63x PL APO NA=1.4 objective (Zeiss) at 37°C and 5% $CO_2$ in an incubator chamber (Incubator PM S1, PeCon, Erbach, Germany). Zen software (Zeiss) was used to define regions of interest along cell membranes of two adjacent transfected cells. The 488 nm laser line was used at 100% transmission to bleach the GFP-Dsg3 signal and fluorescence recovery was recorded over time. ImageJ (NIH) and FrapBot (frapbot.kohze.com) (Kohze et al., 2017) were used for data analysis.

**Image Processing and Statistics**

Figures were compiled with Photoshop CC and Illustrator CC (Adobe, San José, CA, USA). Statistical analysis was performed using GraphPad Prism 8 with a two-tailed student's *t-test* for the comparison of two data sets and one-way or two-way ANOVA corrected by either Dunett's or Tukey's test for more than two data sets. Statistical significance was determined at $p<0.05$. Data shown are mean ± standard error of the mean.

#### SUPPLEMENTARY FIGURES

##### Supp. Fig. S1

(**a**) Left panel: Immunostainings of HaCaT Pg ctrl and ko cell lines stained for Pg. Right panel: Immunostainings of HaCaT Dp ctrl and ko cell lines stained for Dp. DAPI was used in all cell lines to counterstain nuclei. n=3. Scale bar: 20 µm. (**b**) Western blot analysis of adherens junction proteins in Dp as well as Pg ctrl and ko cell lysates. GAPDH served as a loading control. Ctrl pool is the mean value of the three ctrl cell lines. Each ko clone was normalized to the respective ctrl pool, n=3, error bars represent mean ± SEM.

##### Supp. Fig. S2

AFM measurements in a second pair of Dp ctrl and ko cells with (**a-d**) Dsc3 or (**e-h**) Dsg3-coated cantilevers to analyze (**a, e**) distribution of unbinding events in adhesion maps, with each blue pixel representing one unbinding event; green dotted lines mark cell borders, (**b, f**) number of unbinding events, (**c, g**) unbinding force and (**d, h**) the relative interaction probability, classified into bent and plateau unbinding events. Each data point represents the mean value of one biological replicate with >225 analyzed force-distance curves, n≥3, error bars represent mean ± SEM. Scale bar: 2 µm.

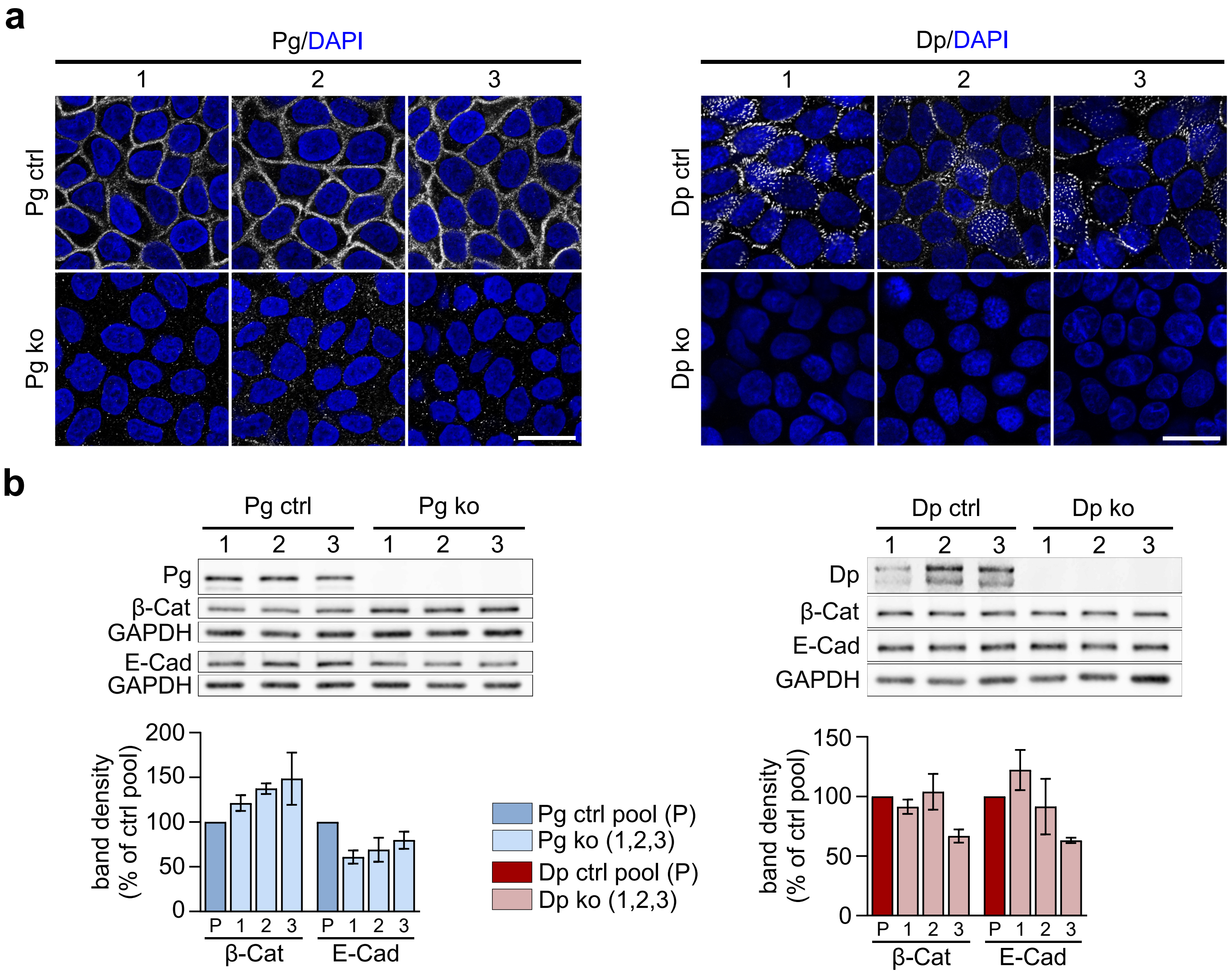


Supplementary Figure S1

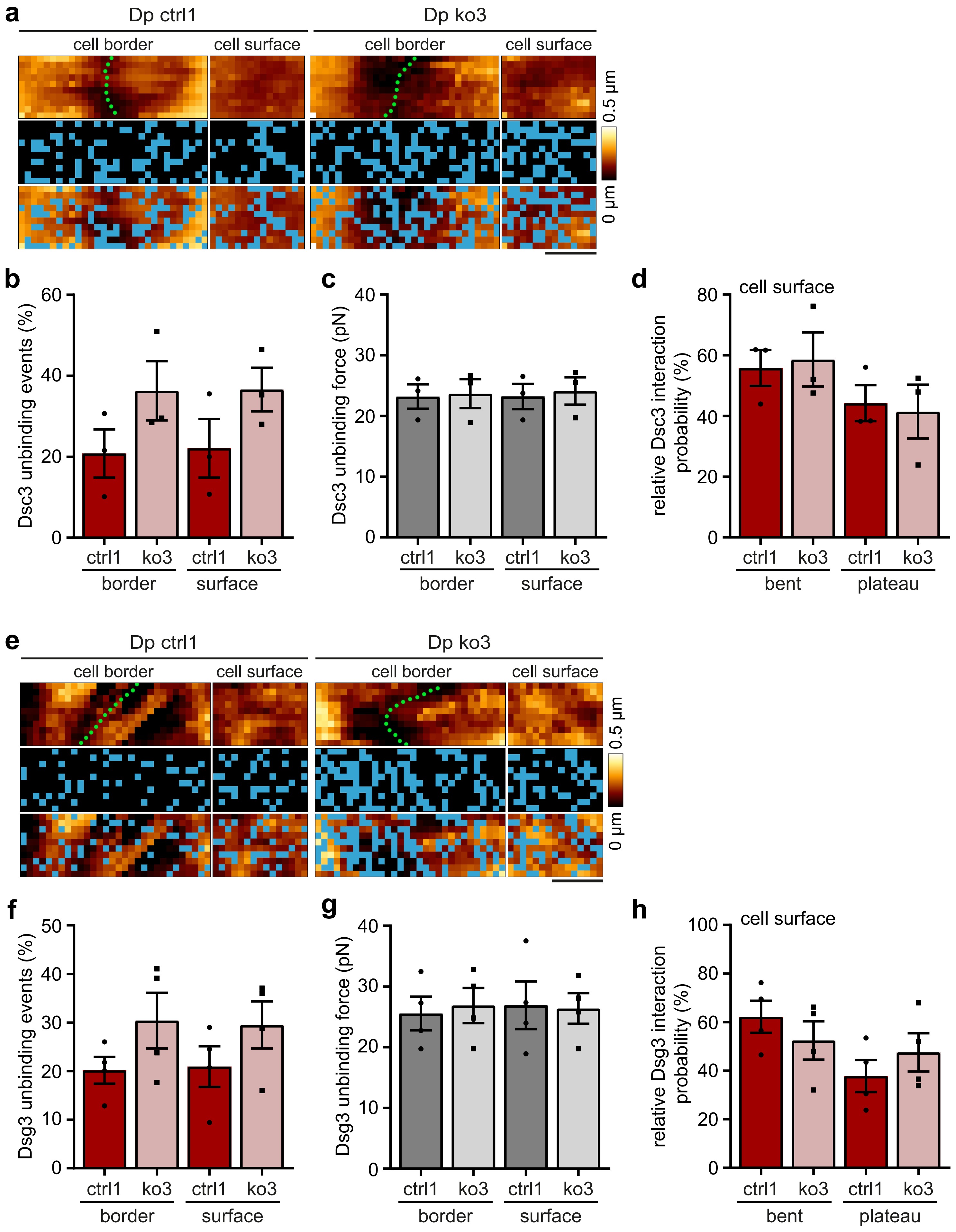


Supplementary Figure S2
